# Key to identification of starch grains used as foods

**DOI:** 10.1101/392209

**Authors:** Artur Manoel Leite Medeiros, Carlos Alexandre Marques

## Abstract

The description of starch grains is important in different areas, such as bromatological, taxonomic, or archaeobotanic analyzes, among others. Although there are data on the microscopic description of starches used as foods, this work has proposed the development of a key to identify the starch grains in 13 raw materials obtained in specialized stores, markets and fairs to facilitate its identification. For optical microscopy, the material was deposited on a slide, being sealed with a coverslip in 50% glycerin. The samples were also mounted on double-sided tape and colloidal graphite on aluminum stubs and analyzed in scanning electron microscopy (SEM). Therefore, an unpublished starch identification key is proposed, together with the update of important data for its morphological description.

## Introduction

Starches are products of the polymerization of glucose mainly in plants during the photosynthesis, responsible for the energy reserve, consisting of grains of different shapes, sizes and stratifications [5, 6, 15].

The description and identification of starch constitutes the basic knowledge to develop diverse investigative research, since its use for analysis of falsifications and adulterations in flours and foods [11, 12], even to the development of hypotheses about the cultivation and domestication of plants in populations of hominids as old as the Paleolithic [2,7].

Although there are data on the microscopic description of starches used as foods, sometimes the inconclusive description of the morphological patterns of very similar starch grains, as well as the lack of detail inherent in clear-chamber drawings, can lead to wrong identifications. However, there is still little research that has like objectives organizes this information in a practical way to the researcher [11, 12, 13]. The authors of these papers use illustrated schemes to characterize the starch grains found in flours. In turn, this ilustrations was based on studies dating from the late nineteenth and early twentieth century [9]. According to them, the microscopic visualization of a starch grain may reveal a single or crossed central or eccentric point or fissure, these formations being called hilum. The authors consider important characteristics to identify starches: the shape, the structure, the state of aggregation and the type of hilum.

Other works [3, 8, 12, 14, 16] present the morphological characterization of starch, but this does not occur in such an embracing way, in the morphological aspect. In addition, there is a relevant study that classified families according to the morphology of the starch [4]. And it is important to mention the identification key to the Mexican Starch Groups, where the most important characters for the classification were the diameter of the starch grains [17].

Therefore, it is necessary a work that organize this informations in a practical way for asecure identification of starch grains. It is relevant to use an identification key, which allows toinform which starch was found in a certain sample in a practical and simple way [17]. Thus,investigative work for starch grains identification can be developed either in bromatology,systematics or archaeobotany, with the use of this tool.

So this paper has proposed to develop a key to identify the starch grains in 13 raw materialsobtained in specialized stores, markets and fairs: oats, rye, barley, corn, wheat, soybeans, beans,bananas, rice, sweet potatoes, false-arrowroot, cassava and potato. The materials was colected in thecities of Rio de Janeiro, Nilópolis and Nova Iguaçu, RJ, Brazil It were also accompanied by thereview and updating of the descriptions and terms found in literature, as well as the registrationthrough photomicrographs and electromicrographs of the material composing the key, to make itpossible for any analyst to rapidly identify such materials.

## Material and Methods

In this paper, 13 raw material were used, from which starch grains were obtained, called: oats (*Avena sativa* L. — Poaceae), rye (*Secale cereale* L. — Poaceae), barley (*Hordeum vulgare* L. — Poaceae), corn (*Zea mays* L. — Poaceae), wheat (*Triticum sp.* L. — Poaceae), soybean (*Glycine max* Merr. — Fabaceae), bean (*Phaseolus vulgaris* L. — Fabaceae), banana (*Musa × paradisiaca* L. — Musaceae), rice (*Oryza sativa* L. — Poaceae), sweet potato (*Ipomoea batatas* Lam. — Convolvulaceae), false-arrowroot (*Maranta ruiziana* Körn — Marantaceae), cassava (*Manihot esculenta* Crantz — Euphorbiaceae) and potato (*Solanum tuberosum* L. — Solanaceae) (Table 1).

**Table 1.** Characteristics of the samples used

| Common name | Flour | Plant organ | Scientific name | Family |
| --- | --- | --- | --- | --- |
| Sweet potato | No | Tuberous root | *Ipomoea batatas* | Convolvulaceae |
| Cassava | Yes | Tuberous root | *Manihot esculenta* | Euphorbiaceae |
| Bean | No | Seed | *Phaseolus vulgaris* | Fabaceae |
| Soybean | No | Seed | *Glycine max* | Fabaceae |
| False-arrowroot | Yes | Rhizome | *Maranta ruiziana* | Marantaceae |
| Banana | No | Fruit | *Musa × paradisiaca* | Musaceae |
| Barley | Yes | Caryopsis | *Hordeum vulgare* | Poaceae |
| Corn | Yes | Caryopsis | *Zea mays* | Poaceae |
| Oats | Yes | Caryopsis | *Avena sativa* | Poaceae |
| Rice | No | Caryopsis | *Oryza sativa* | Poaceae |
| Rye | Yes | Caryopsis | *Secale cereale* | Poaceae |
| Wheat | Yes | Caryopsis | *Triticum sp.* | Poaceae |
| Potato | No | Tuber | *Solanum tuberosum* | Solanaceae |

For analysis in light field and polarization microscopy, the samples were mounted between microscope slide and cover slip, using 50% glycerin (with distilled water) as the mounting medium. To obtain the seed starch, such as beans and soybeans, there was cold extraction in water, after imbibition of the seed for 12 hours, followed by maceration in grail and pistil. In the banana, the technique of direct crushing was made between the microscope slide and cover slip, and in sweet potato and potato, a free-hand cross-section was made [10, 11].

The microscope slides were analyzed using the trinocular photonic microscope, with planchromatic objective, model Alltion ABM103, containing PZO ocular micrometric ruler of 100 μm ± 1 μm. The polarization microscopy was performed using filters for polarization PZO in the same microscope [1].

Micrographs were made by means of an 8MP digital still camera coupled to the photonic microscope and the computer.

For Scanning Electron Microscope (SEM) analysis, the samples were mounted on double tape on 12.7 mm diameter aluminum stubs, both of the Ted Pella Inc., and then analyzed for SEM Phenom World, Pro X model, with load-reducing sample port, no sample metallization required. The electron coils was configured in Image and Point modes at 10 kV and 15 kV, respectively, for generating electromicrographs, at varying microscopic magnifications. The measurements were made by the software contained in the device [1].

## Results and Discussion

In this study, a high frequency of spherical, polyhedral, truncated and ovoid grains was observed. However, the presence of campanulate and round-triangular shaped grains were specific for identification of cassava and false-arrowroot, respectively.

Differences in some classifications were observed, comparing with the bibliography [6, 9, 11, 12]. In sweet potato, truncated grains are reported because their polygonal faces are visible under light microscopy. This is because the authors preferred to name two types of grains: truncated indefinitely and polyhedral. However, a classification that distinguishes indefinitely truncated and polyhedral grains is difficult, which make the characterization become redundant. For this reason, it is preferable was to denominate these grains as polyhedral. However, there is an important subdivision for the truncated grains: the round-triangular grains, which are grains truncated on two faces forming a triangular structure with a spherical/rounded base, being peculiar in false-arrowroot.

The authors also do not cite clearly the presence of spherical and polyhedral grains in cassava, however much their illustrations represent their presence. In wheat, barley and rye, the authors added the presence of ellipsoid grains to describe lenticular grains that revolve around their axis, making their thinner face visible. However, it is preferable to describe as ellipsoids only the predominantly biconvex grains, without the angle of rotation interfering significantly in the classification, since this two-dimensional view is unique to optical microscopy. Potato and banana starches are examples of true ellipsoid grains. In barley, are not cited the in-Y hilum, or any analogous characterization. Even if this characteristic being differential between rye and barley.

It is important to note that barley, rye and wheat have lenticular and spherical grains, which may indicate a peculiar characteristic of Triticeae, the tribe of these vegetables. The same happens with beans and soybean, two species of the tribe Phaseoleae that have spherical, ovoid and reniform grains.

Therefore, the granule descriptions were in accordance with the literature, updating dubious characteristics previously proposed:

- As to state of aggregation: simple or isolated and compound, the compounds subdivided into grouped, aggregated and homogeneously compound;
- As to shape: campanulate, ellipsoid, lenticular, ovoid, polyhedral, pyriform, reniform, round-triangular, spherical and truncated;
- As to the type of hilum: in-Y, linear, punctuated or dotted and star-shaped;
- And as to position of the hilum: concentric and eccentric.

After observing the morphology of the grains already mentioned, the following identification key was made:

## Conclusions

In these paper it was possible to review the knowledge regarding the identification of the most common starches in foods. The identification key shows its relevance and application in several areas with interest in the subject, especially in research involving bromatology and taxonomy. With this, it was possible to update the information described in the literature. Important characteristics for the identification of starch grains, which were redundant or misunderstood in the literature, were updated, and light field microscopy images were confirmed by scanning electron microscopy.

**Figure 1.**
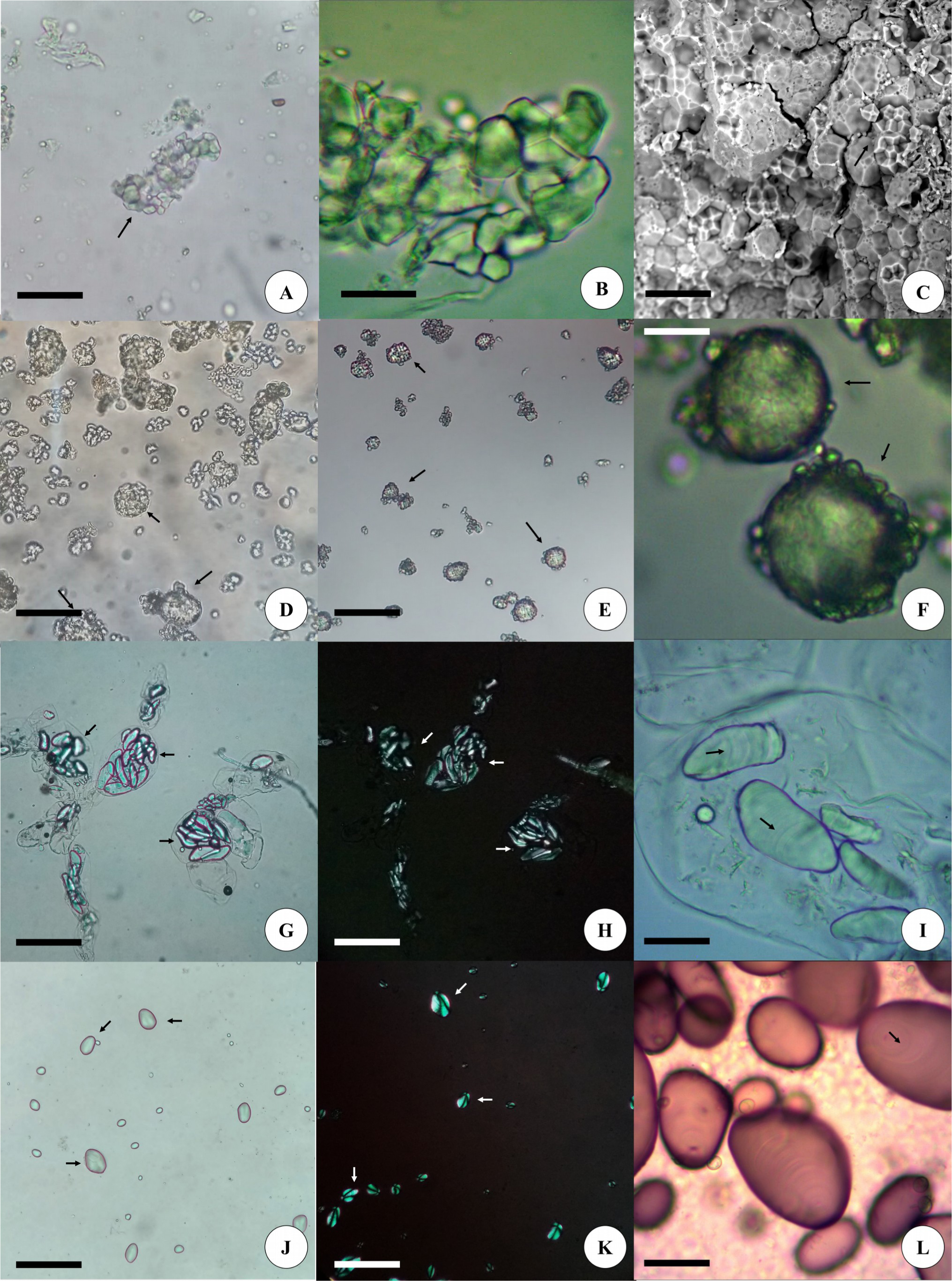
A-C. Rice starch (*O. sativa*). A. grains aggregated in the center of the image (arrow). Bar = 40 μm. B. Aggregate grains in detail. Bar = 20 μm. C. Detail seen in SEM, where the multifaceted aspect of the grains composing an aggregate (arrow) is observed. Bar = 20 μm. D-F. Oat starch (*A. sativa*). D. Homogeneously compound grains (arrows). Bar = 80 μm. E. Characteristic birefringence of homogeneously compound grains (arrows) in polarized light. Bar = 100 μm. F. Detail of the grains (arrows). Bar = 20 μm. G-I. Banana starch (*Musa × paradisiaca*). G. Ellipsoid grains (arrows) within the parthenocarpic fruit pulp parenchyma cells. Bar = 100 μm. H. Characteristic birefringence of banana starch (arrows) in polarized light. Bar = 100 μm. I. Larger grains, allowing the observation of the lamellae (arrows). Bar = 20 μm. J-L. Potato starch (*S. tuberosum*). J. Characteristic pyriform grains (arrows). Bar = 100 μm. K. Extinction cross (well visible) on the grains in polarized light (arrows). Bar = 100 μm. L. grains in detail, sometimes allowing the observation of lamellae (arrow). Bar = 20 μm.

**Figure 2.**
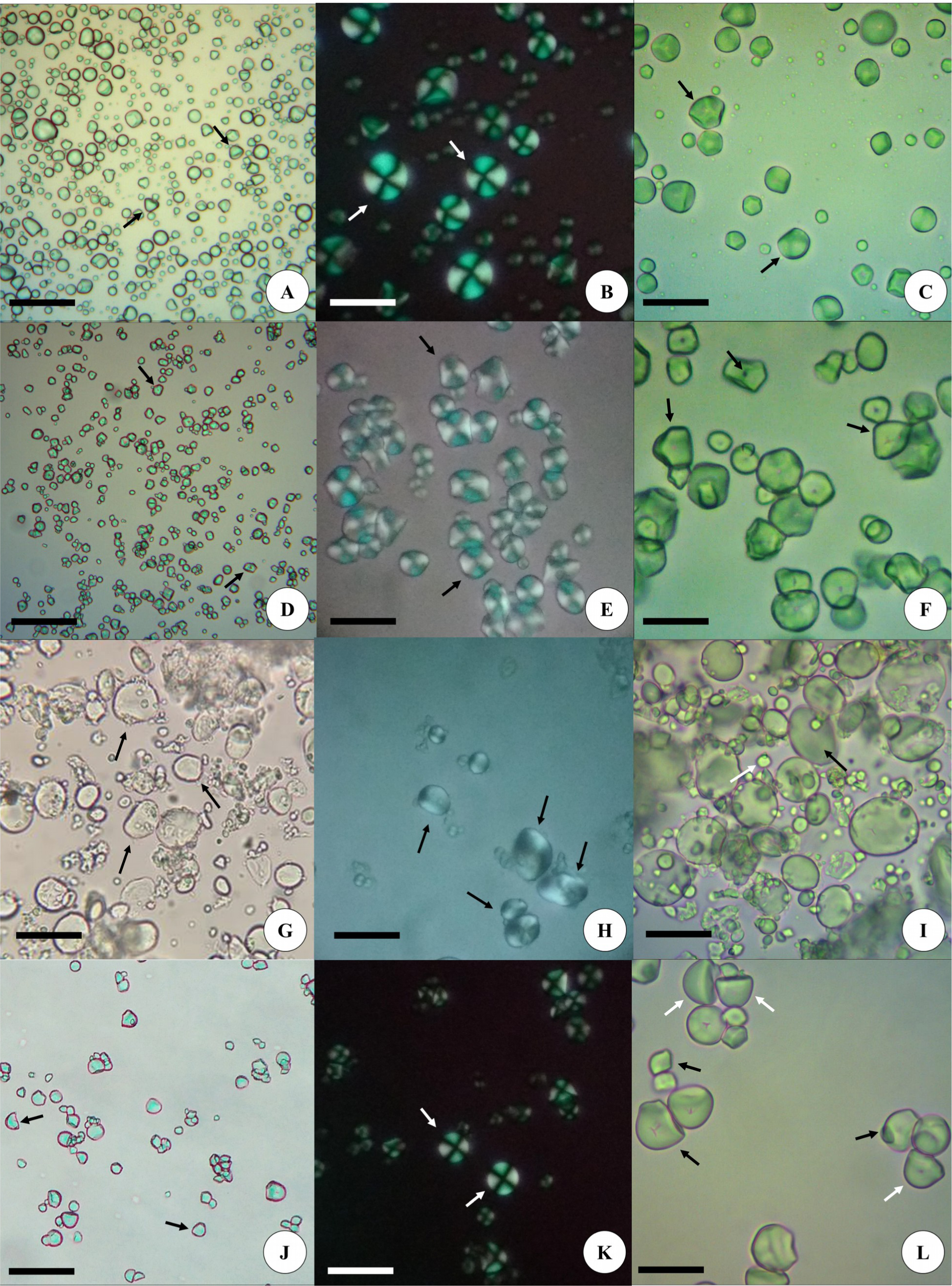
A-C. Sweet potato starch (*I. potatoes*). A. Characteristic campanulate starch grains (arrows). Bar = 100 μm. B. Extinction cross well visible in starches, with characteristic sinuosity (arrows) in polarized light. Bar = 40 μm. C. Predominant polyhedral grains, with presence of campanulate grains (arrows). Bar = 100 μm. D-F. Corn starch (*Z. mays*). D. Typical polyhedral grains (arrows). Bar = 100 μm. E. Highly visible extinction cross (arrows) in polarized light. Bar = 40 μm. F. Polyhedral grains in detail (arrows), showing the geometric aspect of well-marked faces with punctuate hilum. Bar = 20 μm. G-I. Wheat starch (*Triticum sp.*). G. Lenticular grains (arrows) characteristic of the species. Bar = 40 μm. H. Characteristic birefringence of wheat (arrows) in polarized light. Bar = 40 μm. I. Lenticular grains (black arrows) and spherical grains in detail (white arrows). Bar = 30 μm. J-L. Cassava starch (*M. esculenta*). J. Predominant and other truncated polyhedral grains (arrows). Bar = 100 μm. K. Extinction cross well visible in starches (arrows) in polarized light. Bar = 40 μm. L. Grouped campanulate (white arrows) and truncated (black arrows) grains in detail. Bar = 20 μm.

**Figure 3.**
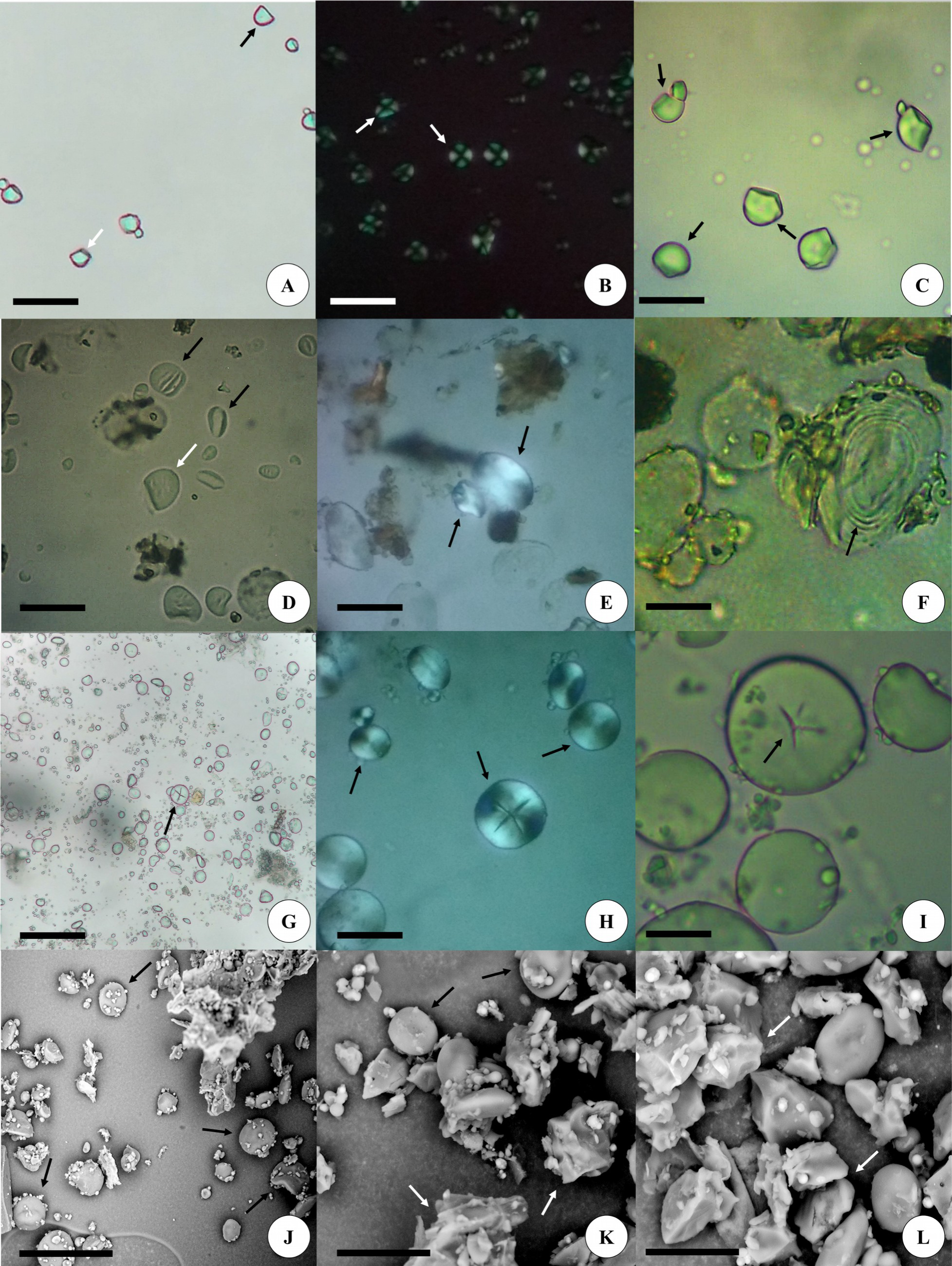
A-C. False-arrowroot starch (*M. ruiziana*). A. Predominant polyhedral grains, others truncated (black arrow) and round-triangular (white arrow). Bar = 100 μm. B. Extinction cross visible in starches (arrows) in polarized light. Bar = 40 μm. Fig. C. Larger round-triangular grains (arrows). Bar = 20 μm. D-F. Barley starch (*H. vulgare*). D. Lenticular grains (black arrows) and truncated grains (white arrow). Bar = 40 μm. E. Characteristic birefringence of barley (arrows). Bar = 40 μm. F. Larger lenticular grains, with detail for the concentric lamellae (arrow). Bar = 20 μm. Fig. G-J. Rye starch (*S. cereale*). Fig. G-H. Lenticular grains, observe the cruciform hilum (arrow). Bar = 40 μm. I. Lenticular grains in greater microscopic magnification, with detail for in-Y grain (arrow). Bar = 20 μm. J. Predominant lenticular grains (arrows) to the SEM. Bar = 80 μm. K-L. Barley starch in SEM. K. Lenticular grains (black arrows) and polyhedral grains (white arrows). Bar = 20 μm. L. Starch grains in detail, highlighting the polyhedral grains (arrows). Bar = 20 μm.

**Figure 4.**
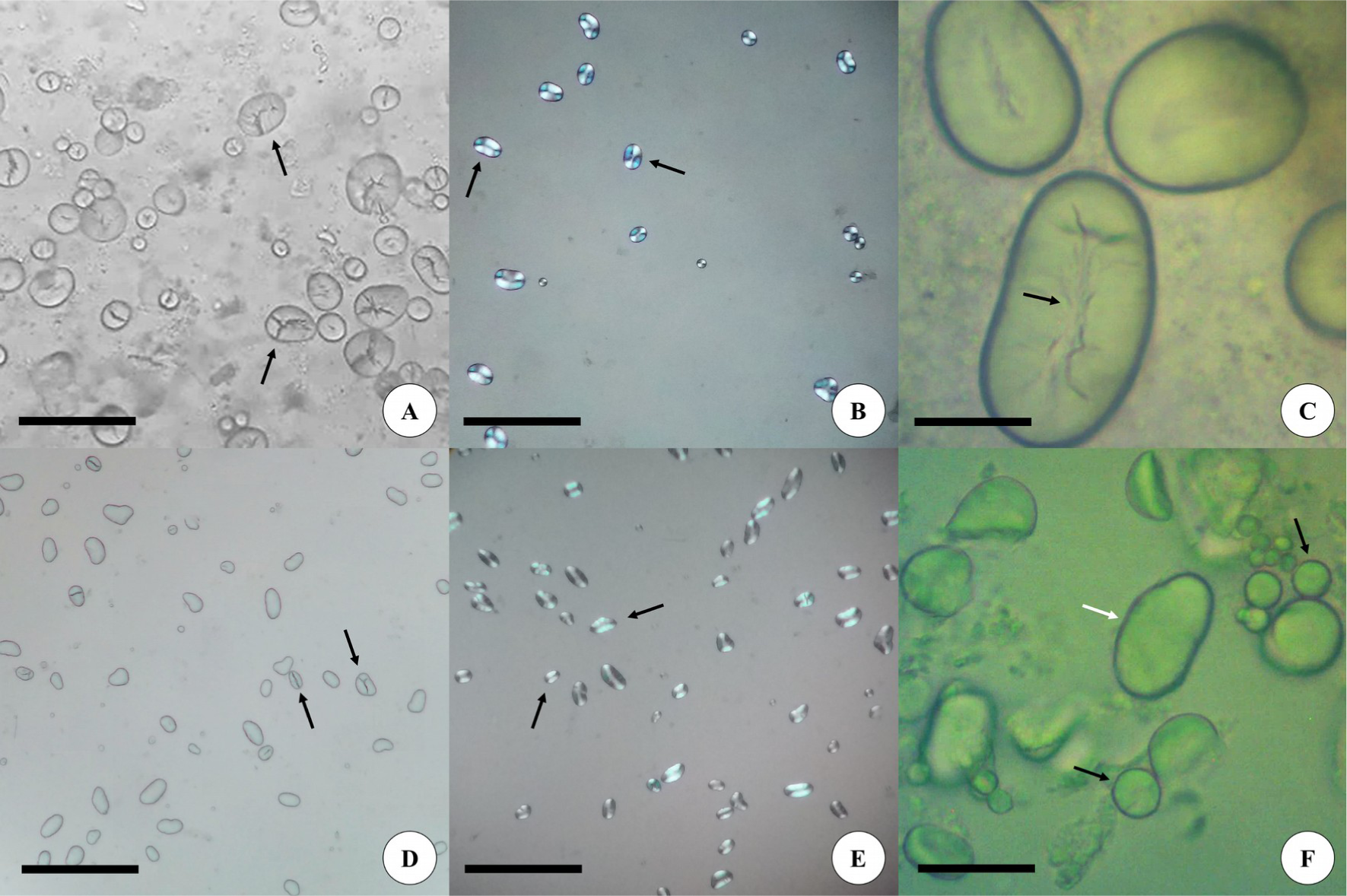
A-C Bean starch (*P. vulgaris*). A. Predominant reniform grains, with characteristic hilum with fissures or lateral branches (arrows). Bar = 80 μm. B. Typical birefringence of the bean, which evidenced its hilum (arrow) in polarized light. Bar = 100 μm. C. Reniform grains in greatest increase. Observe the linear hilum that branches laterally (arrow). Bar = 20 μm. Fig. D-F. Soy starch (*G. max*). D. Predominant reniform grains, with linear hilum (arrows). Bar = 100 μm. E. Typical birefringence of soybean (arrows) in polarized light. Bar = 100 μm. F. Detail where, in addition to the reniform granule (white arrow), spherical and ovoid grains (black arrows) are observed. Bar = 30 μm.

